# Maternal behavioral compensation after neonatal separation fails to prevent spinal circuit reprogramming in offspring

**DOI:** 10.64898/2026.07.03.736384

**Authors:** Hannah Illouz, Apolline Poli, Yasmine Brik, Vincent Lelievre, Pierrick Poisbeau

## Abstract

Early-life adversity durably alters neural development through complex mother-offspring interactions whose underlying mechanisms remain poorly understood. We investigated how neonatal maternal separation (NMS) affects the large repertoire of maternal behaviors and subsequently influences spinal nociceptive circuit development and pain responses in rat offspring.

Rat dams underwent NMS from postnatal day 2 (P2) to P12, 3h/day, and maternal behaviors were assessed before and after the separation period. These behaviors were compared to those of control (non-separated) dams. Offspring spinal cord and dorsal root ganglia were analyzed at P14 and P24 for several neurotrophic, glutamatergic, and GABAergic gene expression patterns. Offspring nociceptive sensitivity was also assessed at P24.

NMS induced increased maternal behaviors (including longer arched-back nursing, higher nest occupancy, and better pup retrieval efficiency), alongside reduced self-care behaviors. These behavioral adaptations were correlated with spinal gene reprogramming in offspring, characterized by a biphasic developmental pattern. At P14, we observed elevated neurotrophic signaling alongside increased GABAergic and glutamatergic markers. By P24, neurotrophic factors decreased while compensatory changes emerged, yet persistent excitatory-inhibitory imbalances remained evident. Parallel to these results, NMS rats also showed mechanical and thermal hot hypersensitivity at P24.

These findings reveal that despite apparent maternal behavioral compensation following NMS, offspring exhibit neurotrophic-driven developmental dysregulation resulting in persistent spinal circuit alterations. The disconnect between maternal behavioral normalization and sustained molecular changes suggests that early separation stress triggers enduring neurobiological cascades independent of ongoing maternal care quantity, with long-term consequences for sensory processing and pain sensitivity.

**Graphical abstract:** 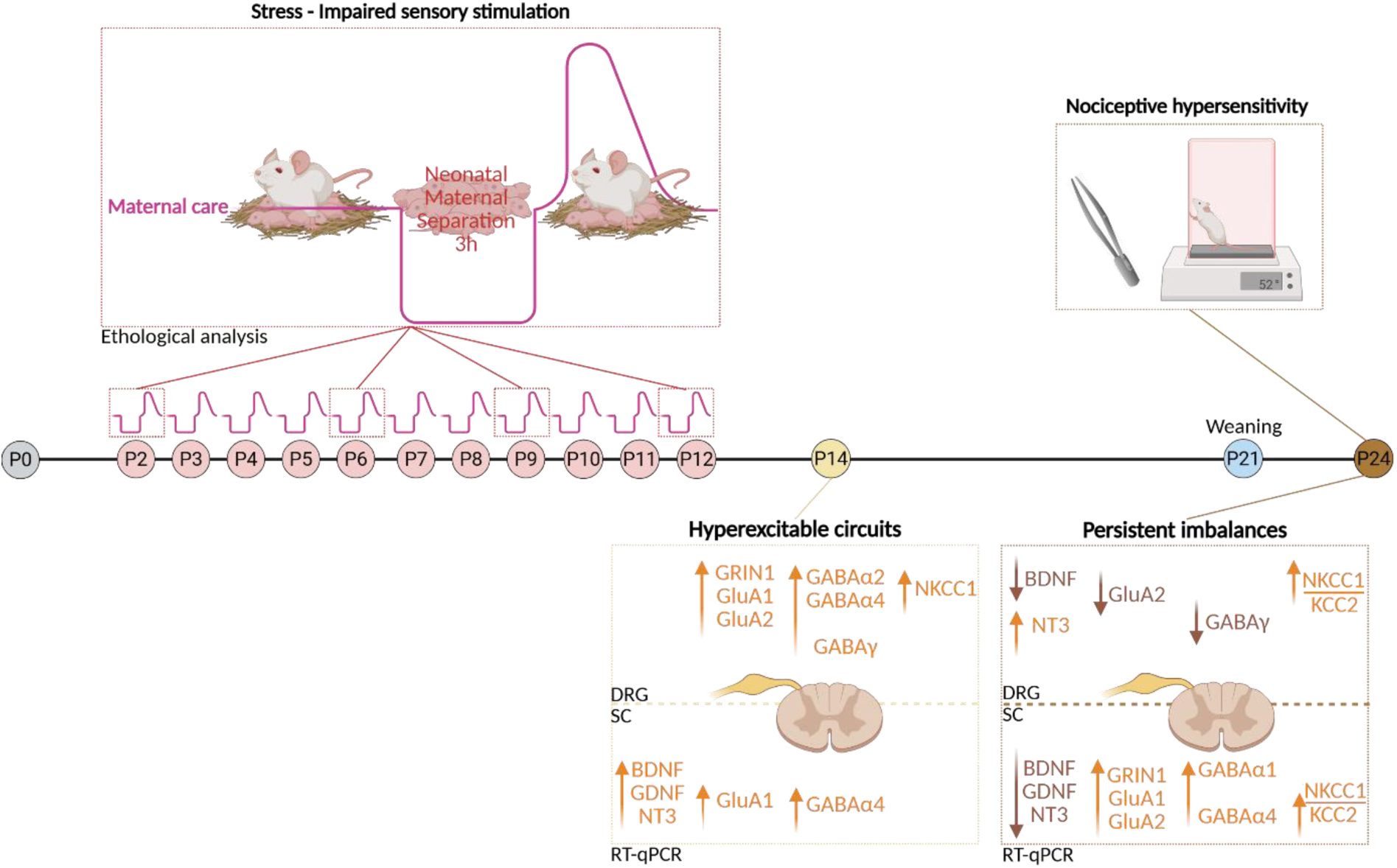

## Introduction

The developmental origins of health and disease (DOHaD) concept proposes that pre-and perinatal experiences, as well as environmental influences, could durably affect gene expression and alter tissue structure and function, thus leading to increased risks of developing various chronic diseases (Barker, 1994; Gluckman and Hanson, 2006). The early postnatal period is of critical importance for brain development, as the nervous system undergoes intense neurodevelopmental processes facilitating proper cognitive, affective, and social development (Moore et al., 2017; Indrio et al., 2023). However, it also renders the brain particularly vulnerable to environmental influences such as early-life stresses (ELS). A robust correlation between childhood exposure to abuse or household dysfunction (adverse childhood experiences, ACEs) and the presence of multiple risk factors for a range of adult mortality causes has already been documented in several studies (Felitti et al., 1998; Pechtel and Pizzagalli, 2011; Herzog and Schmahl, 2018). These ACEs have deleterious consequences on the development of the brain and on later adult behavior, including cognitive and emotional disorders as well as blunted responses to pain, such as increased pain rating or incidence of chronic pain at adulthood (Melchior et al., 2022).

Neonatal maternal separation (NMS) and other rodent models of ELS are helpful paradigms for analyzing the neurobiological substrate responsible for these consequences. NMS consists of isolating the pups from the dam in the early postnatal stage (postnatal day 2 (P2) to P12, 3h/day). This results in long-term sensory, emotional and cognitive consequences. Animals that have been separated present persistent somatic and visceral hypersensitivity (Coutinho et al., 2002; Melchior et al., 2018; Gieré et al., 2023), hypothalamic–pituitary–adrenal (HPA) axis hyperreactivity (Plotsky and Meaney, 1993), increased anxiety (Plotsky and Meaney, 1993; Kalinichev et al., 2000), sociability deficits and mnesic deficits in adulthood (Gazzo et al., 2021; Illouz et al., 2025). To offset these consequences in humans, several studies have focused on early skin-to-skin contact, and kangaroo care clinical studies have documented the long-term benefits of early parental presence and sensory contact (Charpak et al., 1997; Johnston et al., 2017). Studies on the early developmental environment of rodents also showed that it is shaped by maternal influence, with the mother providing essential elements for offspring survival including nutrition, grooming behaviors, and thermal regulation. In this way, the dam controls the maturation of regulatory systems that will influence subsequent behavioral and physiological responses of its offspring (Hofer, 1994).

Natural variations in maternal care quality significantly influence offspring development (Meaney, 2001; Champagne et al., 2003). Specifically, dams can be classified as exhibiting either high or low levels of licking-grooming and arched-back nursing (LG-ABN), with high-LG-ABN mothers displaying more frequent and longer-duration caregiving behaviors (Meaney, 2001; Champagne et al., 2003). Additionally, variations in maternal self-care behaviors, such as eating, drinking, and resting out of nest, can also be affected by environmental stressors and may influence the quality of maternal care provided to offspring. High-LG-ABN offspring show enhanced maternal care, reduced anxiety, lower stress-induced corticosterone levels, and improved learning performance (Liu et al., 1997, 2000; Francis et al., 1999; Bredy et al., 2003; Pedersen et al., 2011). Cross-fostering experiments demonstrated that females born to low-LG-ABN mothers but reared by high-LG-ABN foster mothers exhibited high-LG-ABN maternal profiles similar to their foster mothers (Francis et al., 1999; Weaver et al., 2004), indicating the acquired, non-genetic nature of these behavioral patterns.

These maternal influences, whether natural or induced by stress, seem to be particularly critical for sensory circuit development. While research on ELS has predominantly focused on supraspinal structures, spinal circuits represent an interesting crossroad target structure for developmental programming. As primary processors of sensory information, spinal cord and dorsal root ganglia (DRG) neurons undergo extensive maturation during the early postnatal period, coinciding with the critical window for maternal care effects. This developmental period extends through the third postnatal week, with the development of the sensory and nociceptive system starting before birth and following a similar neurodevelopmental pattern in the rat and in the human fetus (Verriotis et al., 2016; Melchior et al., 2022). Maternal care could play a crucial role in regulating these developmental processes, as it considerably stimulates the sensory system of the offspring during the first weeks of life. Indeed, imitating the tactile stimulation derived from maternal licking/grooming could partially protect the offspring from behavioral and neuroendocrine consequences of ELS (Lomanowska and Melo, 2016; Kentner et al., 2018). Conversely, prolonged maternal absence notably triggers a cellular shift from growth and development toward conservative survival-focused metabolic maintenance (Fish et al., 2004).

The maturation of these spinal circuits is critically influenced by neurotrophic factors, particularly brain-derived neurotrophic factor (BDNF) and glial cell line-derived neurotrophic factor (GDNF), which serve as key regulators of GABAergic and glutamatergic system development (Boyce and Mendell, 2014). In particular, this developmental process enables the maturation of chloride homeostasis, controlling the switch from depolarizing to hyperpolarizing GABA (Rivera et al., 1999; Cordero-Erausquin et al., 2005; Brewer et al., 2020). This phenomenon has been proposed to play a critical role in the central sensitization of nociceptive circuits (Poisbeau and Salvat, 2025). ELS interferes with this maturational process, as it decreases the expression of BDNF expression (Roth et al., 2009; Boersma et al., 2014), and NMS delays this GABA excitatory-to-inhibitory functional switch (Furukawa et al., 2017). In addition, variations in maternal-offspring interactions might permanently alter the subunit composition of the GABAA receptor complex in the offspring, which may explain the relationship between early life events and vulnerability for anxiety disorders (Caldji et al., 2003).

Despite these documented consequences on offspring, the mechanisms by which NMS disrupts the maternal-offspring interactions remain poorly understood. It is well documented that dams subjected to NMS exhibit elevated plasma corticosterone levels, as well as modified hypothalamic neuropeptide expression patterns associated with stress and anxiety responses (Maniam and Morris, 2010). These dams also display distress behaviors towards their pups beginning as early as P6, indicating that NMS constitute a stressful experience for the dams, likely modifying maternal behavior. Several studies show that variations in maternal caregiving behaviors when dams are reunited with their pups can explain individual differences in adult offspring’s HPA axis reactivity (Denenberg, 1999; Huot et al., 2004). In fact, pups that receive more maternal stimulation have lower ACTH, corticosterone, and CRH mRNA, as well as greater amounts of GR mRNA (Liu et al., 1997; Denenberg, 1999). Moreover, when dams receive foster litters during separation periods, the biological pups do not develop HPA axis-hyperreactivity in adulthood (Huot et al., 2004), indicating that NMS consequences are partially attributable to the stress-inducing effects of the separation on the mother herself.

Given that ELS outcomes often resemble those observed in offspring of low-LG-ABN mothers, there is a tendency to assume that ELS models necessarily result in reduced maternal care quality and quantity (Burenkova and Grigorenko, 2024). Research examining maternal behavioral adaptations during NMS has yielded conflicting findings. Some studies demonstrate increased LG-ABN and nest occupancy during post-separation periods, suggesting compensatory responses (Biggio et al., 2014; Couto-Pereira et al., 2016), while others report decreased maternal behaviors or no differences compared to controls (Romeo et al., 2003; Der-Avakian and Markou, 2010). The methodological heterogeneity across studies, including inconsistent reporting of essential maternal behaviors and varying observation timepoints, complicates direct comparisons and highlights critical gaps in our understanding of how maternal behavioral adaptations during NMS may influence offspring neurodevelopmental outcomes (Orso et al., 2019).

While maternal behavioral adaptations to separation stress have been partially documented, their influence on offspring spinal circuit maturation remains largely unexplored. Moreover, no study has linked maternal caregiving during the NMS period to specific molecular changes in offspring spinal circuits. This study aims to investigate maternal behavioral adaptations during NMS and their potential impact on offspring spinal circuit development. To address this, maternal care was systematically and finely assessed between P2-P12. Spinal gene expression patterns were analyzed at P14 and P24, and nociceptive thresholds assessed at P24. This approach aims to establish temporal relationships between maternal behavioral responses to NMS and the emergence of persistent spinal circuit alterations in offspring, providing mechanistic insights into how ELS programs long-term pain vulnerability.

## Materials and Methods

### Animals

Pregnant female Sprague-Dawley rats (Charles River, Saint-Germain Nuelles, France) were used in this study and carefully monitored for delivery. The day of birth was referred to as postnatalday 0 (P0). Mother and pups were housed in a temperature (22 ± 2 ◦C) and humidity (45 ± 10 %) controlled room, under a 12-hour light–dark cycle (lights on at 7:00 am), with ad libitum access to food and tap water. Pups were weaned at P21 and housed in collective cages according to sex. Males and females were used in this study, and results were pooled as no sex-specific difference was observed. Different litters were used for molecular analyses (2 litters/group). All procedures were conducted in accordance with EU regulations and approved by the regional ethical committee (CREMEAS authorization numbers APAFIS #34227-2021112914341095 v9).

### Neonatal maternal separation

Litters were randomized at birth into two groups: nonseparated control (CTRL) litters and NMS. From postnatal days 2 (P2) to 12 (P12), the litters assigned to the NMS group were taken out of the nest cages three hours a day and put in a different cage under a heating lamp (28 ◦C wavelengths between 1400 nm and 3000nm – one cage/litter). During the whole NMS time, the litters assigned to the CTRL group stayed in their home cages with their mothers and did not get any additional treatment aside from changing their cage bedding, as well as maternal care assessments. The pups were weaned at P21 and kept in cages with four rats each.

### Maternal care assessments

#### Pup retrieval test

Retrieval behavior was assessed during postpartum on days 2, 6, 9 and 12 after delivery. In the CTRL group the litter was separated from the dam for less than 2 min and immediately put back in the homecage. In NMS groups, this test was performed immediately after the separation period. The pups were placed into one corner of the cage opposite to the nest site. The following behaviors were scored: latency to first pup-contact (latency to sniff or contact the first pup); 2 pup-retrieval (latencies to retrieve 2 pups back to the nest), whole-litter-retrieval (latencies to retrieve the litter back to the nest), number of pups in nest at the end of test. The test ended after 15 min, or when the female had retrieved all the pups.

#### Nest building observations

As homecages were changed (P7/P8), old nesting material was removed and 2 new nesting material blocks were put in each new cage. The nest quality was rated on a 5-point scale using a naturalistic nest scoring system from Hess *et al*., 2008. The scores range from 0 (undisturbed nesting material), 1 (evident interaction with nesting material), 2 (flat nest), 3 (cup nest), 4 (incomplete dome) and 5 (complete dome). Scoring occurred 1-, 24- and 48-hours after cages were changed.

#### Behavioral observations

Observations of the dams were carried out in two filming sessions per day at postnatal days 2, 6, 9 and 12. Behavioral observation filming sessions lasted 30min and took place in the morning between 9:30 and 11:30 and in the afternoon between 13:30 and 15:30, when white lights were on and animals were in their inactive phase. In the NMS groups, the afternoon film occurred directly after the pup retrieval test, e.g. after the separation period when mother and offspring were reunited. Behavioral analysis of the videos was then performed on the basis of previously employed ethological parameters (Chourbaji et al., 2011). Behaviors comprised: ‘licking/grooming’, ‘active nursing’, ‘passive nursing’ and ‘nest building’ as manifestations of caring behavior (Sup.fig.1); ‘self-grooming’ and ‘eating/drinking’ as a manifestation of dam self-maintenance (Sup.fig.2); and ‘dam out of nest’ as a behavior that related neither to the dam’s immediate self-care nor her interacting with her litter (Sup.fig.3). During the analysis, each of the mother’s behaviors was timed minute by minute over the 30-minute period. Different behaviors could be timed at the same moment if they coexisted (e.g., out of nest self-grooming). All videos were analyzed blind (neither the corresponding group, nor the day, nor the timing of the video was known to the experimenter during viewing and analysis).

### Molecular analyses

#### Tissue processing

The lumbar segment of the spinal cord (SC), as well as dorsal root ganglions, were collected at P14 or P24 with n=7/group. SC and DRG were harvested and stored at 80 ◦C.

#### RNA extraction and RT-qPCR analysis

Total RNA was extracted using a protocol adapted from the original procedure of Chomczynski & Sacchi, 1987 consisting in 2 independent total RNA extractions separated by a DNAse I treatment (TURBOTM DNase; Ambion, Life technologies, Saint Aubin, France), as previously described in detail (Lelievre et al., 2002). 800 ng RNA were reverse transcripted with the RT iScript kit (Bio-Rad, Marnes-la-Coquette, France). Quantitative PCR was performed using SYBR Green Supermix (Bio-Rad), on the iQ5 Real Time PCR System (Bio-Rad). Amplifications were carried out in 42 cycles (20 s at 95 ◦C, 20 s at 60 ◦C, and 20 s at 72 ◦C). Primer sets for all genes of interest were designed using Oligo6.0 and M fold softwares (primer sequences in Supplementary Table 1). Samples were accurately dispensed in duplicates using a robotic workstation (Freedom EVO100; Tecan, Lyon, France), and amplification efficacy given by standard curves was always close to 100% (±2%), while amplification specificity was assessed by a melting curve study. Data were normalized to the housekeeping hypoxanthine–guanine phosphoribosyltransferase (HPRT) since its transcripts remained highly stable among the different samples and analyzed using the ΔΔct method (Livak and Schmittgen, 2001).

### Nociceptive assessments

#### Mechanical nociception

Mechanical nociception was measured with a calibrated forceps (Bioseb, Vitrolles, France) as previously described (Luis-Delgado et al., 2006). The habituated rat was loosely restrained with a towel masking the eyes to limit stress by environmental stimulations. The tips of the forceps were placed at each side of the hindpaw, and a gradually increasing force was applied. The pressure producing withdrawal of the paw corresponded to the nociceptive threshold value.

#### Thermal nociception

Thermal nociception was assessed using hot- and cold-plate tests (Hot-Cold Plate, Bioseb, France). Without prior habituation, animals were placed in a Plexiglas compartment with a hot or cold plate at the bottom, maintained at 52 ◦C or 0 ◦C respectively. The latency of onset of the first nociceptive behavior (licking or withdrawal of one or more paws, hobbling, or jumping) was noted and corresponds to the rat’s thermal nociceptive threshold to hot or cold. If no nociceptive behavior is observed after 60 s, the animal is removed from the chamber to avoid potential tissue damage.

#### Statistical analyses

Maternal care behavioral data were analyzed using linear mixed-effects models (LMM) with Group (CTRL-NMS), Day (2-6-9-12), and Timing (before-after) as fixed effects and individual dam as a random effect. Residual normality was assessed using Shapiro-Wilk tests. When normality assumptions were violated (p < 0.05), data were log- or square-root-transformed to stabilize variance and improve normality. If transformations were insufficient, generalized linear mixed models (GLMM) with beta distribution were used. When GLMM failed to converge due to sparse data structure, we retained LMM on sqrt-transformed data as a conservative approach. This method was preferred over non-parametric alternatives given our complex repeated-measures design with random effects, which cannot be adequately handled by distribution-free tests. Type II or III analysis of variance (2w or 3wANOVA) was performed on the selected model for each behavior. Significant main effects and interactions (p < 0.05) were followed by post-hoc pairwise comparisons using estimated marginal means (emmeans) with Tukey adjustment for multiple comparisons.

Unpaired t-tests were used to compare mechanical and thermal nociception between CTRL and NMS rats, after residual normality was verified using Shapiro-Wilk tests.

All PCR results were processed using the ΔΔct method (Livak and Schmittgen, 2001). To account for the distinct biological nature of DRG and spinal cord tissues, RT-qPCR results were assessed with separate 2wANOVAs (GroupxTime) for each region followed by Sidak’s multiple comparison post-hoc test.

Maternal care behavioral data analyses were conducted in R (version 4.5.1) using the lme4, lmerTest, glmmTMB, and emmeans packages. Analysis and graphical representation of transcript expression and nociceptive behavioral data was performed using GraphPad Prism 9 software (La Jolla, USA).

For each experiment, normality of residuals and homoscedasticity was verified with Shapiro-Wilk and Spearman’s tests. Differences were considered statistically significant for p < 0.05. Data are expressed as mean ± standard error of the mean (SEM).

## Results

### NMS dams exhibit higher motivation and better efficacy to care for their offspring after 3h-separation

Maternal motivation was assessed using repeated pup retrieval tests (at P2, P6, P9 and P12, fig 1A1, 1A2, 1A3), as well as a nest building test scored at P7, P8 and P9 (fig 1B).

The LMM analysis on log-transformed data of the pup retrieval test revealed that NMS dams took significantly less time to have the first contact with their pups (fig.1A1; main effect of Group, F(1,54.94)=9.30, p=0.004; estimated marginal mean (EMM) difference = 0.83 log-seconds, SE=0.28), suggesting greater motivation and potentially indicative of increased anxiety. Significant changes across postpartum period were also observed (main effect of day, F(3, 52.17)=3.42, p=0.024), with day 2 being significantly faster than day 12 (t(52.17)=2.71, p=0.043). No significant effect of the interaction between Group and Day was measured (F(3, 52.17)=0.55, p=0.649). Similarly, NMS dams took less time to retrieve 2 pups back in nest, which indicates increased efficacy (fig.1A2). This latency significantly decreased between P2 and P6, suggesting that dams learned and became familiar with the test (main effect of group, F(1,54.07)=5.13, p=0.027; EMM difference = 0.44, SE=0.20 and main effect of day, F(3, 52.19)=3.27, p=0.028; P2-P6: t(52.19)=2.87, p=0.029; no effect of GroupxDay). However, there were no significant differences in the latency to bring back the entire litter (fig.1A3), either in comparisons between groups (F(1,53.82)=0.04, p=0.852) or over the course of days (F(3, 51.58)=1.42, p=0.247), which can be explained by the fact that several mothers (CTRL and NMS) did not bring the entire litter back to the nest before starting to nurse the pups (15 minutes cut-off latency).

LMM analysis on untransformed data of the nest building score of dams on day 7, 8 and 9 (1, 24 and 48 hours after cages were changed) were in accordance with pup retrieval results as it revealed that NMS mothers reconstructed the nest around their litter more efficiently than CTRL mothers (fig. 1B; main effect of Group, F(1, 37.25)=11.93, p=0.001; EMM difference = −0.67 points, SE=0.19), evoking increased motivation. For both groups, the score increased over time (main effect of Hours, F(2, 35.70)=74.77, p<0.0001) with a clear difference between 1 hour after the change and 24 (t(35.70)=-9.96, p<0.0001) or 48 (t(35.70)=-11.02, p<0.0001) hours after, indicating rapid nest rebuilding. No significant effect of the interaction between Group and Hours was measured (F(2, 35.71)=1.68, p=0.200).

**Figure 1:**
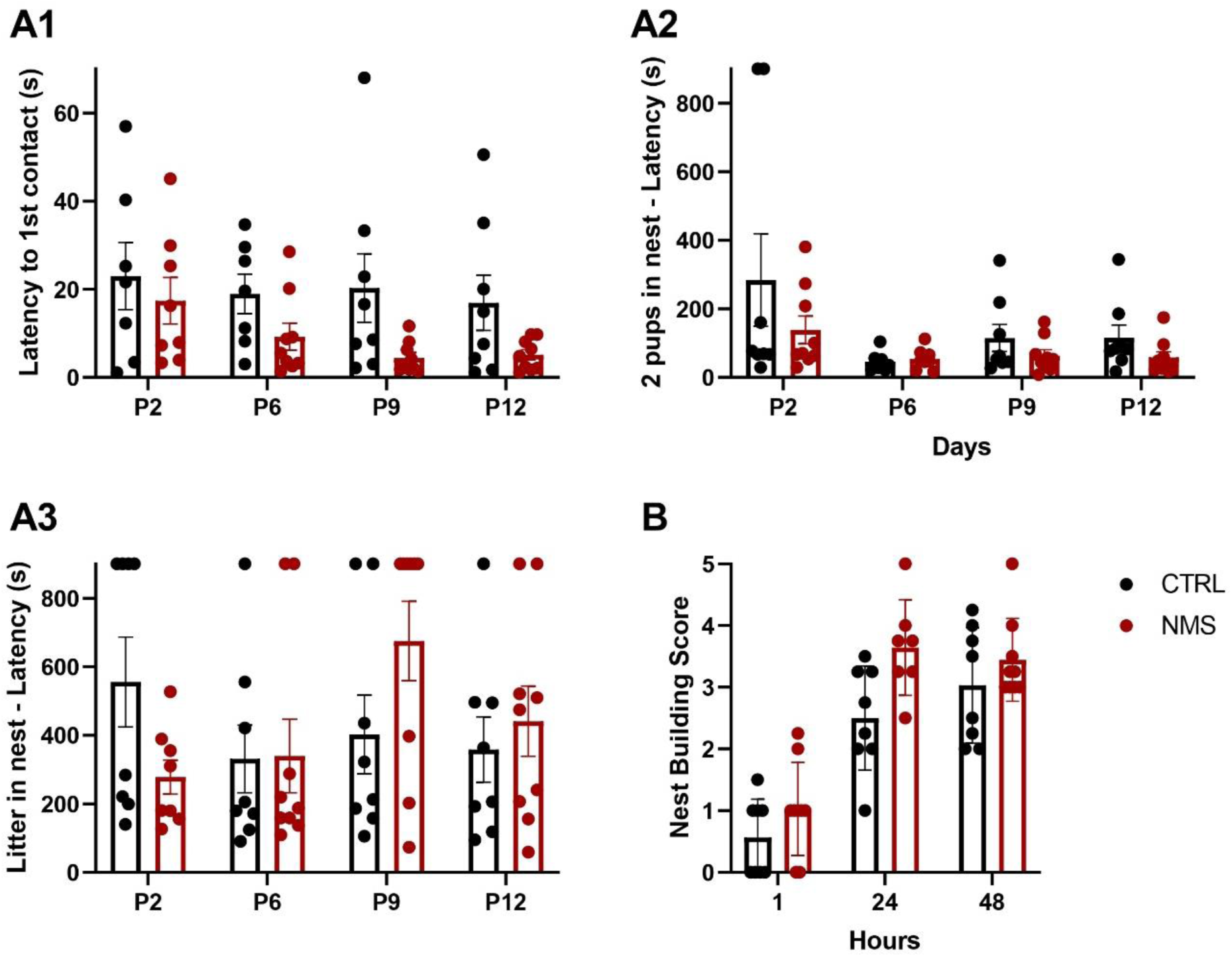
Maternal motivation assessed with pup retrieval test (A) and nest building test (B). Time until first contact between the dam and their pups (A1), as well as the latency to retrieve 2 pups (A2) and the whole litter in nest (A3) were timed. The nest building score (B) completes this maternal motivation analysis. Statistical significance was assessed using LMM on log-transformed data (A1, A2, A3) or on untransformed data (B). 2wANOVA was performed on the selected model, followed by post-hoc pairwise comparisons using estimated marginal means (emmeans) with Tukey adjustment for multiple comparisons. This analysis revealed a main effect of Group and Day concerning latency to first contact (A1) and the latency to bring 2 pups back in nest (A2), as well as a main effect of Group and Hours concerning nest building scores (B). CTRL: N = 8, NMS: N = 9.

### Neonatal maternal separation increases the duration of active and pup-directed behaviors while reducing passive and dam-directed behaviors

Behavioral analysis before and after the three hours of separation revealed specific behaviors particularly affected by NMS. At first glance, the duration of nursing behaviors (active and passive), as well as the moments when the dam was inside or outside the nest, were the highest and varied most between the two groups (fig. 2A). The heat map also indicated that most variations seemed to happen directly after (and not before) the separation period.

**Figure 2:**
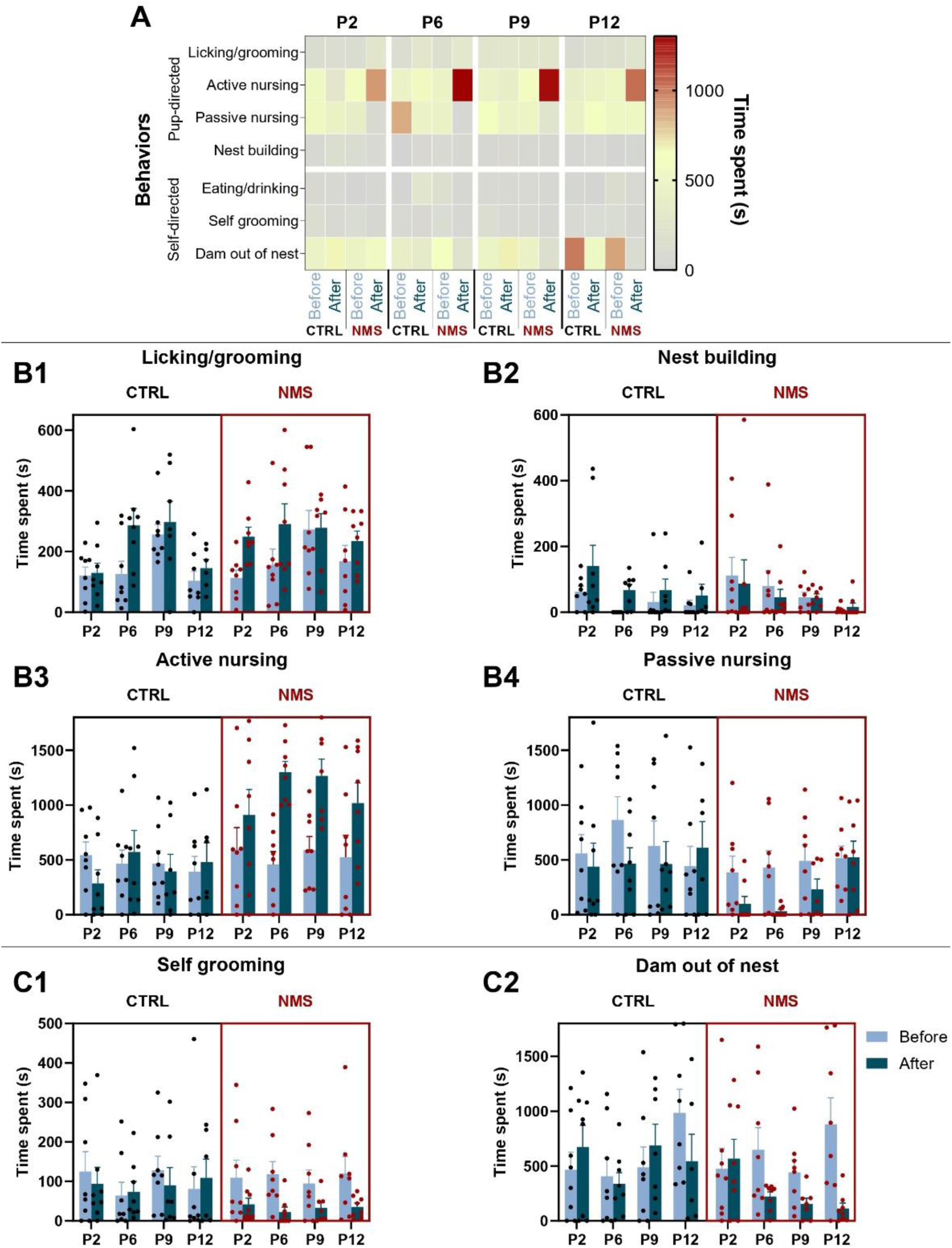
Effects of NMS on the range and time allocation of maternal behaviors. Each dam was observed for 30min and 7 behaviors were timed before or after pup retrieval test (control group) or NMS. The heatmap (A) enables identification of the most recurrent behaviors. 4 behaviors are pup-directed (B), while 3 behaviors are centered on the dam (self-directed, C). Statistical analysis was performed using LMM on square-root-transformed data (B1, B3, C2) or on log-transformed data (B2, B4, C1). Statistical significance was assessed with 3wANOVA, followed by post-hoc pairwise comparisons using emmeans with Tukey adjustment for multiple comparisons. A **main effect of Group** was revealed concerning active and passive nursing (B3, B4). Licking/grooming presents a **main effect of Day and Timing**. Nest building (B2), Active nursing (B3), Self-grooming (C1) and Dam out of nest (C2) display an **interaction effect of GroupxTiming**. CTRL: N = 8, NMS: N = 8.

Licking and grooming (LG) were the behaviors that varied least between groups (fig. 2B1). LMM analysis on square-root transformed data indicated no significant main effect of Group, but an upward trend in the duration of LG in the NMS group (F(1, 101.47)=3.09, p=0.082). A significant effect of Day and Timing was observed (main effect of Day, F(3, 101.44)=5.31, p=0.002) and main effect of Timing, F(1, 101.29)=12.12, p=0.0007), with a specific increase between P2 and P9 (t(101.44)=-3.57, p=0.003) and decrease between P9 and P12 (t(101.44)=3.30, p=0.007). An increase in LG after the separation period was also specifically noticeable (EMM difference = −2.96, SE=0.85). LMM analysis on log-transformed duration of nest building (NB) showed no main difference between CTRL and NMS groups (F(1, 108)=0.04, p=0.846) but a specific decrease in duration across days (main effect of Day, F(3, 108)=3.50, p=0.018, P2-P12: t(102)=3.13, p=0.012), as the pups grew up (fig. 2B2). Additionally, an interaction effect between Group and Timing highlighted a difference between CTRL mothers, which rebuilded their nest extensively after the pup retrieval test, and NMS mothers, which did not do so after the separation period (F(1, 108)=8.60, p=0.004, EMM difference = −1.72 log-seconds, SE=0.50). Arising from these results, a tendency toward an effect of Timing (F(1, 108)=3.89, p=0.051) and GroupxDay interaction (F(3, 108)=2.38, p=0.074) has also been revealed.

Active nursing (AN, fig.2B3) is a specific type of nursing during which the mother presents a recognizable, energy-demanding arched-back posture. LMM analysis on square-root data revealed that NMS mothers spent significantly more time actively nursing their litters (main effect of Group, F(1,108)=16.79, p<0.0001, EMM difference = −7.98, SE=1.95). The main effect of Timing (F(1,108)=6.29, p=0.014, EMM difference = −4.89, SE=1.95) and the interaction effect between Groups and Timing (F(1,108)=12.35, p=0.0006, NMS before-NMS after: t(101)=-4.30, p<0.0001) also pointed out that this increase is specific to the period following the 3 hours of separation. No significant effects of Day (F(3, 108) = 0.83, p = 0.480) or GroupxDay were observed (F(3, 108) = 0.22, p = 0.882), indicative of a stable effect over time.

Passive nursing (PN, fig.2B4) behavior analysis showed results that complement those of AN. In fact, when AN increased, it appeared that PN decreased (fig.2A). LMM on sqrt-transformed data revealed a significant lower duration of PN from NMS mothers compared to CTRL (main effect of Group, F(1,108)=6.12, p=0.015, EMM difference = 5.45, SE=2.2), as well as a decrease after the separation period compared to before (main effect of Timing, F(1,108)=6.49, p=0.012, EMM difference = 5.61, SE=2.2). These results were the exact opposite of AN, suggesting that the previously described increase in AN came at the expense of PN. No significant effects of Day (F(3,108) = 0.96, p = 0.417) or interaction effects were observed (all p > 0.05).

In general, the accumulation of behaviors directed toward pups (licking/grooming, nursing, nest building; fig.2A) increased among NMS mothers (LMM analysis on sqrt-transformed durations, main effect of Group, F(1,108)=4.26, p=0.041, EMM differences = - 3.24, SE=1.57); particularly just after NMS (Interaction effect GroupxTiming, F(1,108)=5.30, p=0.023, NMS before-NMS after: t(101)=-2.87, p=0.005). On the other hand, behaviors directed toward the mother herself (self-directed: eating/drinking, self-grooming, out of nest; fig.2A) decreased significantly after separation in NMS mothers (LMM analysis on sqrt-transformed durations, main effect of Timing, F(101.3)=4.25, p=0.042, EMM difference = 4.43, SE=2.15); Interaction effect GroupxTiming, F(1,101.5)=6.38, p=0.013, NMS before-NMS after: t(101.5)=9.87, p=0.001). This effect was particularly pronounced on day 12, following pups-development over time (Interaction effect TimingxDay, F(3,101.5)=2.74, p=0.047, Day12 before-Day12 after: t(102)=3.15, p=0.002).

As the time measurement intervals were 30 minutes, few mothers drank or ate during this period, which did not allow for an optimal analysis of this behavior (Sup.fig.4). However, it appeared that CTRL mothers spent more time eating or drinking specifically in the period before the pup retrieval test, compared to after (LMM analysis on sqrt-transformed durations, Interaction effect GroupxTiming, F(1,101.5)=5.39, p=0.022, NMS before-NMS after: t(101)=-1.73, p=0.087). Concerning self-grooming (fig.2C1), NMS mothers spent significantly less time in this behavior following the 3-hour separation (LMM analysis on sqrt-transformed durations, main effect of timing, F(1,101.2)=5.06, p=0.027, EMM difference=2.11, SE=0.94; Interaction effect GroupxTiming, F(1,101.3)=5.43, p=0.022, NMS before-NMS after: t(101)=3.27, p=0.0015), consistent with the general trend of self-centered behaviors mentioned earlier. The same applies to the time NMS dams spent away from the nest after separation (fig.2C2): it decreased significantly (LMM analysis on sqrt-transformed durations, main effect of timing, F(1,101)=4.25, p=0.042, EMM difference=4.33, SE=2.1; Interaction effect GroupxTiming, F(1,101.5)=4.34, p=0.040, NMS before-NMS after: t(101)=2.96, p=0.004). In addition, the time spent away from the nest at P12 is much higher before separation (Interaction effect DayxTiming, F(3,101.5)=2.75, p=0.047, P12 before-P12 after: t(102)=3.26, p=0.001), highlighting the fact that the time spent away from the nest by mothers increased over time in both groups.

### NMS induces changes in neurotrophic factor transcripts expression, and disrupt chloride homeostasis in the spinal cord

Given that early stress has been shown to reprogram nociceptive and sensory circuit development through alterations in neurotrophic factors, chloride homeostasis, and ion channel expression (Juif et al., 2016), the molecular development of offspring sensory-spinal networks was investigated at critical timepoints to determine whether compensatory maternal behaviors observed in the NMS model similarly affect circuit maturation.

Fig. 3A1 indicates that BDNF mRNAs in the DRG were 4 times less expressed in NMS animals compared to CTRL rats, specifically at P24 (2wANOVA, interaction effect TimexGroup, F(1,24)=6.73, p=0.016, CTRL P24 – NMS P24: t(24)=2.98, p=0.013). In the SC, an opposite expression pattern was observed between P14 and P24 in NMS rats: a 3 times BDNF mRNAs overexpression at P14 and a 4 times underexpression at P24 compared to CTRL rats was detected (2wANOVA, interaction effect TimexGroup, F(1,24)=17.59, p=0.0003, CTRL P14 – NMS P14: t(24)=2.6, p=0.031, CTRL P24 – NMS P24: t(24)=3.33, p=0.0056). The relative DRG GDNF expression was unchanged between CTRL and NMS rats (fig. 3A2, 2wANOVA, No significant effects of Time: F(1,24)=0.68, p=0.417; Group: F(1,24)=0.002, p=0.962); or interaction effects: F(1,24)=0.68, p=0.417). The same expression pattern as BDNF was observed in the SC of NMS rats: a 3.3 times GDNF mRNA overexpression at P14 and a 8.5 times underexpression at P24 compared to CTRL rats (2wANOVA, interaction effect TimexGroup, F(1,24)=26, p<0.0001, CTRL P14 – NMS P14: t(24)=2.59, p=0.031, CTRL P24 – NMS P24: t(24)=4.62, p=0.0002). Concerning NT3, a significant mRNA overexpression in NMS DRG was revealed (fig. 3A3, 2wANOVA, main effect of Group, F(1,23)=16.43, p=0.0005, predicted means difference = −1.575, SE=0.39). In the same way as for BNDF and GDNF, an 4.3 times increased NT3 mRNA expression was measured at P14 in the SC of NMS animals, while its mRNA expression is 3.5 times decreased at P24 compared to CTRL (2wANOVA, interaction effect TimexGroup, F(1,24)=18.52, p=0.0002, CTRL P14 – NMS P14: t(24)=3.28, p=0.006, CTRL P24 – NMS P24: t(24)=2.81, p=0.019). NT4 DRG mRNA expression was unchanged between CTRL and NMS rats (fig. 3A4, 2wANOVA, No significant effects of Time: F(1,24)=0.01, p = 0.906; Group: F(1,24)=0.85, p=0.365; or interaction effects: F(1,24)=0.01, p = 0.906). To a lesser extent, the same spinal expression pattern as other neurotrophins was noted (2wANOVA, interaction effect TimexGroup, F(1,24)=5.007, p=0.035). These results highlight a coherent pattern in the developing SC of NMS rats, with an increased expression at P14 and a decreased mRNA expression at P24 compared to CTRL.

**Figure 3:**
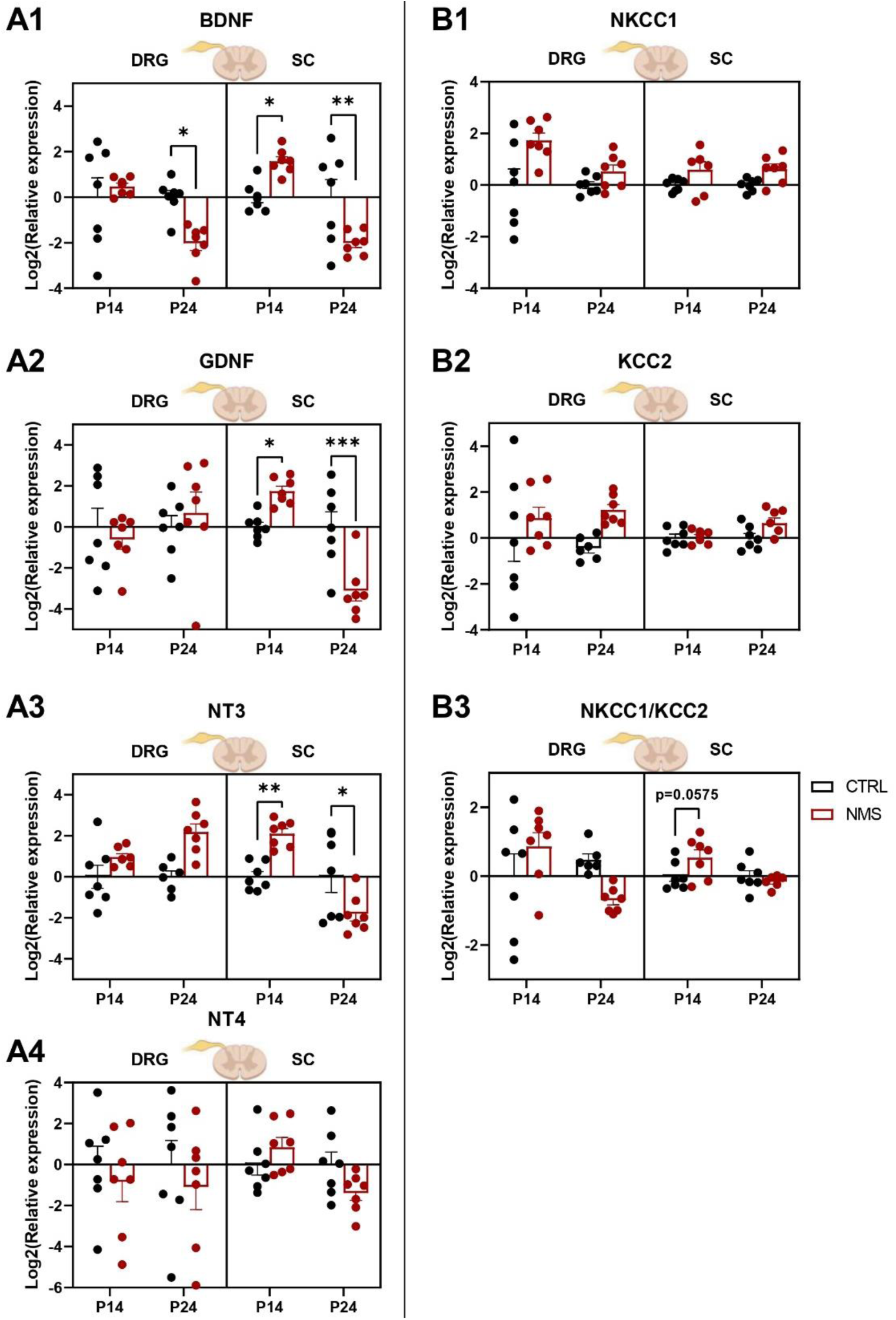
Expression of neurotrophic factors (A) and chloride cotransporters (B) transcripts in developing DRG and SC following neonatal maternal separation (P14 and P24). Statistical significance was assessed with 2-way ANOVAs or mixed-effect analysis on log2-transformed data; followed by Sidak’s multiple comparison post-hoc test, illustrated as follows: p < 0.05 (*), p < 0.01 (**), p < 0.001 (***). A **main effect of group** was detected for NT3, NKCC1 and KCC2 mRNA expression in the DRG, as well as NKCC1 in the SC. BDNF mRNA transcripts in the DRG, and BDNF, GDNF, NT3, NT4, NKCC1/KCC2 in SC presented a **main effect of time**. An **interaction effect of TimexGroup** was displayed concerning dorsal root ganglia’s BDNF mRNA and NKCC1/KCC2, as well as BDNF, GDNF, NT3, NT4, NKCC1/KCC2 mRNA in the SC. CTRL and NMS at P14 and P24 in DRG and SC: N = 7.

Given that BDNF regulates the developmental expression and plasticity of chloride cotransporters, ensuring an optimal neuronal inhibition at adulthood, we next examined NKCC1 and KCC2 transcript levels.

Fig. 3B1 reveals an overexpression in NKCC1 mRNA expression in the DRG of NMS rats in comparison to CTRL (2wANOVA, main effect of Group, F(1,24)=9.25, p=0.006, means difference = −1.127, SE=0.37). A similar significant overexpression was shown in the SC (2wANOVA, main effect of Group, F(1,24)=9.59, p=0.005, means difference = −0.6028, SE=0.19). In the same way, KCC2 mRNA seemed to be generally overexpressed in the DRG of NMS rats at P14 and P24 (fig. 3B2, 2wANOVA, main effect of Group, F(1,23)=4.38, p=0.0475, difference between predicted means = −1.267, SE=0.6051). However, this was not the case in the SC (No significant effects of Time: F(1,24)=2.77, p = 0.109; Group: F(1,24)=3.78, p=0.064; or interaction effects: F(1,24)=2.77, p = 0.109), resulting in a disrupted NKCC1/KCC2 mRNA balance specifically at P14 in the SC (fig. 3B3, 2wANOVA, main effect of time: F(1,23)=4.33, p=0.049; interaction effect GroupxTime: F(1,23)=4.33, p=0.049, CTRL P14 – NMS P14: t(23)=2.24, p=0.0575). This chloride homeostasis balance also seemed to be modified in the DRG of NMS rats (2wANOVA, interaction effect: F(1,23)=6.22, p = 0.0203).

### Neonatal maternal separation alters the development of glutamatergic and GABAergic receptor subunit expression in the DRG and the SC

Neurotrophic factors and chloride cotransporters during postnatal development is likely to affect the function and expression of ionotropic neurotransmitter receptors that mediate excitatory and inhibitory synaptic transmission. The GluA1 and GluA2 subunits of AMPA receptors, as well as the NMDA receptor subunit GRIN1 display similar fluctuations in NMS compared to CTRL rats. In fact, NMS rats displayed higher mRNA levels at P14 in the DRG, suggesting an early peripheral increase in tetrameric glutamatergic ionotropic receptors (2wANOVA, interaction effect TimexGroup, fig. 4A1 GluA1: F(1,24)=6.07, p=0.021, CTRL P14 – NMS P14: t(24)=2.66, p=0.027; fig. 4A2 GLuA2: F(1,23)=20.63, p=0.0001, CTRL P14 – NMS P14: t(23)=3.98, p=0.001; fig. 4A3 GRIN1: F(1,24)=10.27, p=0.004, CTRL P14 – NMS P14: t(24)=3.76, p=0.002). However, this increase in the DRG did not maintain at P24 (fig. 4A1 and 4A3, GluA1: CTRL P24 – NMS P24: t(24)=0.82, p=0.664; GRIN1: CTRL P24 – NMS P24: t(24)=0.77, p=0.694), while GluA2 mRNA is 2.3 times lower in NMS rats (fig. 4A2, CTRL P24 – NMS P24: t(23)=2.48, p=0.0415). At the spinal level, fig. 4A1 and 4A3 show an overexpression in GluA1 and GRIN1 mRNA expression of NMS rats (2wANOVA, main effect of Group, GluA1:, F(1,24)=22.69, p<0.0001, means difference = −0.709, SE=0.15; GRIN1: F(1,23)=12.78, p=0.002, means difference = −0.589, SE=0.16) and no differences between NMS and CTRL groups concerning GluA2 (fig. 4A2, 2wANOVA, no significant effects of Time: F(1,24)=2.98, p = 0.097; Group: F(1, 24)=3.01, p=0.096 or interaction effects: F(1,24)=2.98, p = 0.097). Thus, an early post-NMS response was observed in the periphery (DRG) and a more sustained response at the central level (SC), suggesting a potential increase in excitatory transmission in nociceptive and sensory circuits.

**Figure 4:**
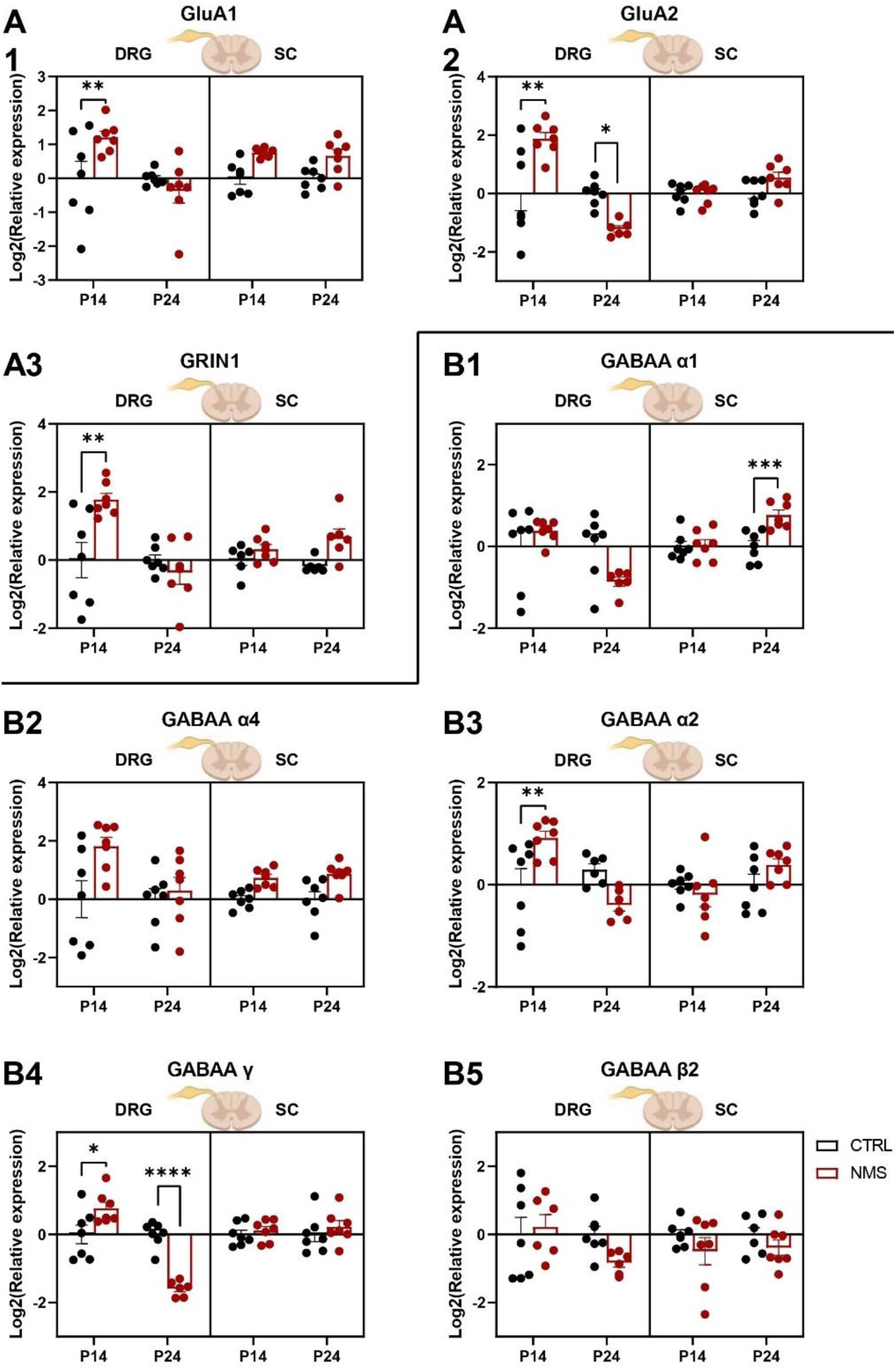
AMPA (A1, A2), NMDA (A3) and GABAA receptor subunit (B) RNA expression in developing sensory circuits of CTRL and NMS rats, as measured with RT-qPCR. Statistical significance was assessed with 2-way ANOVAs or mixed-effect analysis on log2-transformed data; followed by Sidak’s multiple comparison post-hoc test, illustrated as follows: p < 0.05 (*), p < 0.01 (**), p<0.0001 (****). A **main effect of Group** was revealed concerning GRIN1, GABAAα4 and GABAAγ mRNA in the DRG; and GluA1, Grin1, GABAAα1 and GABAAα4 in the spinal cord. GluA1, GluA2 and GRIN1 display a **main effect of time** and an **interaction effect** in the DRG exclusively. GABAAα1, GABAAα2 and GABAAγ present a main effect of time and an interaction effect in the DRG, while GABAAα1 presents these effects also in the spinal cord. CTRL and NMS at P14 and P24 in DRG and SC: N = 7.

In parallel with excitatory glutamate receptors, we examined the expression of some GABAA receptor subunits, as the excitatory/inhibitory balance is critical for proper nociceptive and sensory processing. Fig. 4B1 shows different GABAAα1 mRNA expression levels in the NMS DRG (2wANOVA, interaction effect TimexGroup, F(1,23)=5.68, p=0.026, CTRL P14 – NMS P14: t(23)=1.05, p=0.513, CTRL P24 – NMS P24: t(23)=2.29, p=0.062), as well as in the SC (2wANOVA, interaction effect TimexGroup, F(1,23)=7.81, p=0.01), specifically 1.7 times higher at P24 than in CTRL SC (CTRL P24 – NMS P24: t(24)=4.14, p=0.0007). In the case of GABAAα4 (fig.4B2), this subunit presented a significantly increased expression in the DRG and SC of NMS rats (2wANOVA, main effect of Group, DRG: F(1,24)=5.34, p=0.03, means difference = −1.055, SE=0.46; SC: F(1,24)=21.4, p=0.0001, means difference = −0.8, SE=0.17). GABAAα2 mRNA transcripts were 1.9 times higher in the DRG of NMS rats at P14 (fig. 4B3, 2wANOVA, interaction effect TimexGroup, F(1,22)=15.91, p=0.0006, CTRL P14 – NMS P14: t(22)=3.35, p=0.006), but not different from CTRL in the SC (2wANOVA, no significant effects of Time: F(1, 24)=2.94, p = 0.099; Group: F(1,24)=0.32, p=0.579 or interaction effects: F(1,24)=2.94, p = 0.0997). Significant differences were noticeable concerning GABAAγ mRNA expression in the DRG of NMS animals (fig. 4B4, 2wANOVA, interaction effect TimexGroup, F(1,23)=38.41, p<0.0001): it was 1.7 times higher than CTRL at P14 (CTRL P14 – NMS P14: t(23)=2.91, p=0.016) and 3 times lower than CTRL at P24 (CTRL P24 – NMS P24: t(23)=5.80, p<0.0001). At central level, CTRL and NMS GABAAγ mRNA levels were similar (2wANOVA, no significant effects of Time: F(1,24)=0.11, p=0.738; Group: F(1,24)=1.04, p=0.318 or interaction effects: F(1,24)=0.11, p = 0.738). No difference in GABAAβ2 transcript expression was found in either the DRG or SC of NMS and CTRL rats (fig. 4B5, 2wANOVA, DRG: no significant effects of Time: F(1,22)=2.21, p=0.1515; Group: F(1, 22)=0.75, p=0.395 or interaction effects: F(1,22)=2.21, p=0.1515; SC: no significant effects of Time: F(1,24)=0.04, p=0.841; Group: F(1,24)=2.89, p=0.102 or interaction effects: F(1,24)=0.04, p=0.841). A similar expression pattern was therefore found at the peripheral level: an increase in GABAA subunits at P14, and then a decrease at P24. At the central level, only the levels of GABAAα1 and GABAAα4 subunits increased, particularly at P24, i.e. in the long term.

### Neonatal maternal separation induces mechanical and thermal hot nociceptive hypersensitivity at adolescence

In parallel to molecular alterations shown at P24 (fig. 3 and 4), nociceptive tests revealed that NMS rats exhibited hypersensitivity to both mechanical and thermal hot stimulation after weaning (P25) compared to the control group. They displayed 10.4% lower mean pressure thresholds (fig. 5A; Unpaired Student t test: p<0.0001, t(26)=5.35). NMS animals also showed 28% lower paw withdrawal latencies to noxious heat (fig. 5B; Unpaired Student t test: p=0.0028; t(27)=3.289). However, NMS rats sensitivity to noxious cold was similar to CTRL rats at P25 (fig. 5C; Unpaired Student t test: p=0.747; t(27)=0.326).

**Figure 5:**
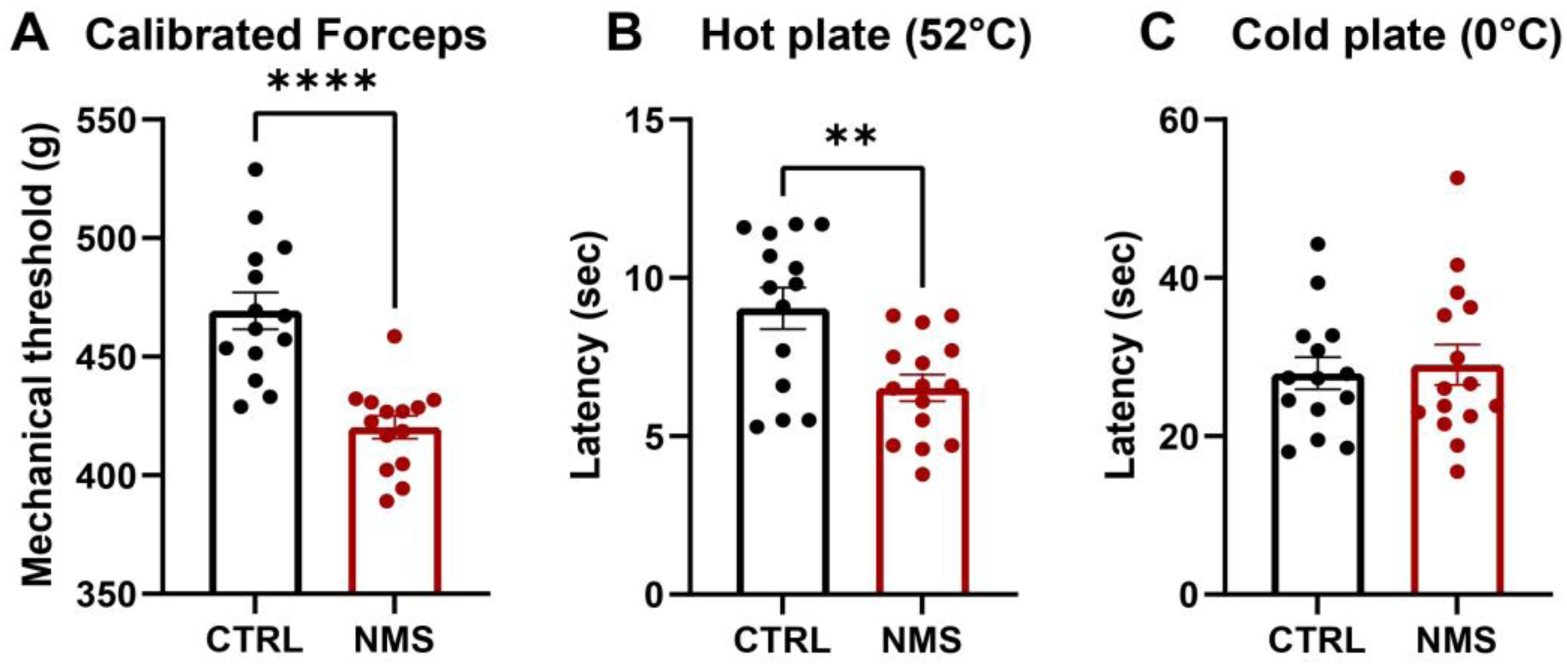
Effect of neonatal maternal separation (NMS) at P25 on mechanical and thermal nociception. NMS induced a significant decrease in mechanical nociceptive thresholds as evaluated with calibrated forceps (A). The hot plate (52 ◦C) test (B) revealed thermal hot hypersensitivity in NMS rats; in contrary to cold plate (0°C) test (C). Statistical significance was assessed with an unpaired t-test, illustrated as follows: p < 0.01 (**), p < 0.0001 (****). CTRL: N = 14, NMS: N = 15.

## Discussion

This study demonstrates that NMS dams exhibit heightened maternal motivation and increased pup-directed care, characterized by faster first contact with pups, more efficient nest building, prolonged active nursing, and reduced self-directed behaviors following the separation period. These behavioral adaptations are associated with molecular changes in the developing somatosensory system of the pups. In the DRG, NMS is associated with an early transient increase in glutamatergic and GABAA receptor subunits mRNA at P14, with most returning to control levels by P24. In the SC, NMS rats present a biphasic pattern of neurotrophic factor expression (elevated mRNA at P14 followed by marked reductions at P24) alongside disrupted chloride homeostasis, and persistent increases in glutamatergic and GABAA receptor subunits. These alterations suggest a shift toward enhanced excitatory transmission and altered inhibitory signaling. Functionally, these molecular perturbations are correlated with nociceptive mechanical and thermal hot hypersensitivity at P25.

Studies establishing a classification into two groups of mothers (high LG-ABN and low LG-ABN; Meaney, 2001; Champagne *et al*., 2003) led to a paradigm suggesting that high LG-ABN is beneficial for offsprings. In fact, naturally high LG-ABN mothers give birth to offspring with reduced stress reactivity, increased hippocampal glucocorticoid receptors expression, enhanced stress resilience, and better cognitive outcomes (Caldji et al., 1998; Champagne et al., 2001; Weaver et al., 2004). A number of research thus associate low LG-ABN and reduced maternal care with ELS, as both induce long-term deleterious consequences for the pups. However, in our study, NMS dams show enhanced maternal activity and motivation, and offsprings still present nociceptive hypersensitivity at adolescence along with increased anxiety and impaired spatial memory (Illouz et al., 2025). These results are only partially supported by the literature, since long maternal separation was mainly considered to reduce maternal care (Caldji et al., 2000; Lovic et al., 2001; Maniam and Morris, 2010; Aguggia et al., 2013; Demarchi et al., 2023), and has only recently been suggested to increase maternal behavior such as LG or ABN (Orso et al., 2019; Baracz et al., 2020; Bölükbas et al., 2020; Demarchi et al., 2023). To our knowledge, our study is the first to take into account most of the parameters mentioned in the literature (Orso et al., 2019) and to focus on behaviors directed toward the dam, in addition to those directed toward the offspring. While previous studies in rats did not really assess the time in or out of nest, the present study shows that NMS dams spend more time in nest compared to CTRL dams, in addition to increased maternal motivation and care. This result parallels the ones from recent research using the limited bedding and nesting model (Gallo et al., 2019), suggesting hypervigilant and stressed mothering, as if the maternal behaviors were anxiety-driven rather than spontaneously nurturing. Indeed, NMS dams show behavioral markers of anxiety in this study, like faster retrieval, intense nest building and sacrifice of self-care; and the literature revealed NMS-associated anxiety in the elevated plus maze, maternal HPA axis hyperactivation during NMS period, as well as increased corticosterone levels in dams (Maniam and Morris, 2010; Alves et al., 2020; Baracz et al., 2020). NMS dams also show increased ultrasonic vocalizations, likely reflecting anxiety-driven hypervigilance (Bölükbas et al., 2020). We hypothesize that, having elevated corticosterone during compensatory care, the dam may transmit stress to pups. Thus, the compensatory care observed in this study may be driven by the dam’s own stress and anxiety state, but could also exist due to pup distress vocalizations, as augmented ultrasonic calling from stressed pup is followed by increased caregiving behavior by the dam immediately upon return to the nest (Bell et al., 1974). Moreover, active nursing increased specifically after separation (not before), suggests a reactive, and not trait-based behavior. The shift from passive to active nursing observed in this study also points out that the reactive care from NMS mothers is an energy-demanding strategy. This could be a reason for resource reallocation, leading to reduced self-directed behaviors and establishing self-sacrifice for offspring care.

Overall, these maternal behaviors create a modified sensory environment with enhanced tactile (nursing), thermal (nest), and chemical (pheromones) stimulation. This leads to strong activation of sensory-spinal pathways, comparable to the effects of certain types of environmental enrichment (Oers et al., 1998; Castelli et al., 2020; Melo and Hoffman, 2024). Notably, the quality of tactile stimulation and nursing during maternal care is well-established as a form of sensory enrichment (Oers et al., 1998; Castelli et al., 2020; Costa et al., 2020; Melo and Hoffman, 2024). However, the compensatory maternal care in the NMS model seems insufficient to counterbalance the cumulative stress of repeated 3-hour separations, particularly if the dam transmits her own elevated corticosterone levels during care delivery. This brings out a critical distinction between stress-induced reactive/compensatory care and naturally occurring high LG-ABN and confirms that compensatory maternal care is not fully able to reverse developmental alterations induced by the dam’s and offspring’s distress experienced during NMS (as also suggested by Wang *et al*., 2020). The offspring outcomes in our study suggest that different molecular programming occurred compared to natural high LG-ABN. This draws on several previous studies demonstrating that ELS induces lasting consequences on certain epigenetic mechanisms: Moloney and colleagues showed that visceral hypersensitivity resulting from this maternal deprivation is associated with reduced histone acetylation in the spinal cord at adulthood (Moloney et al., 2015), while recent results from our team corroborate these findings, as the mechanical nociceptive hypersensitivity classically described in the NMS model was associated with increased expression of certain HDACs, several chromatin methylation proteins, and microRNAs at the spinal level (Illouz et al., 2023).

The differential mRNA expression patterns observed between the DRG and SC provide mechanistic insights into whether the molecular alterations originate peripherally (driven by sensory input changes) or more centrally. Our findings reveal transient molecular changes in the DRG, with early upregulation of certain glutamate receptor subunits and GABAA receptor subunits at P14 that normalize by P24. This pattern could reflect the DRG’s role as the primary target of altered maternal stimulation during the early postnatal period (Fitzgerald, 2005), with initial mRNA upregulation representing an adaptive response to modified sensory input. However, NT3, NKCC1, and KCC2 mRNA remain elevated until P24, suggesting sustained alterations in neurotrophin signaling and chloride cotransporter-mediated homeostasis, specifically at the level of sensory neuron cell bodies in the DRGs. In contrast, neurotrophic factors in the SC display a biphasic pattern, with initial elevation at P14 followed by marked suppression at P24. These early increases in neurotrophins resemble previously published data indicating that litters of high LG-ABN mothers exhibit higher levels of BDNF (Liu et al., 2000). This change is accompanied by disruption of the NKCC1/KCC2 ratio, which appears mechanistically linked to the biphasic BDNF expression. BDNF is a central regulator of spinal development, modulating chloride homeostasis and GABAergic maturation through TrkB-mediated dowregulation of KCC2 (Rivera et al., 2002; Garraway and Huie, 2016), as well as the development of glutamatergic and GABAergic synapses (Gottmann et al., 2009). During development, the ratio of NKCC1 (chloride importer) to KCC2 (chloride exporter) in the SC progressively decreases, enabling the developmental switch of GABA from excitatory to inhibitory (Rivera et al., 1999). However, in NMS offspring, BDNF spike at P14 likely hindered KCC2 expression during this critical window, while the subsequent P24 suppression could not prevent completion of the maturation process, leaving circuits in an immature state: elevated NKCC1 (immature characteristic) with inadequate KCC2 upregulation (incomplete maturation). This maintains an impaired neuronal chloride extrusion, leading to depolarizing GABA and sensitized sensory and nociceptive circuits in the SC at P25. In addition, the specificity of this pain phenotype (mechanical and thermal hot, but not cold hypersensitivity) suggests selective alterations in Aδ and C-fiber pathways rather than global sensory disruption (Basbaum et al., 2009; Gieré et al., 2021), consistent with neurotrophins’ surplus-mediated sensitization of peripheral nociceptors (Fitzgerald, 2005).

The sustained overexpression of glutamate receptor subunits in the SC of NMS animals, combined with altered GABAA receptor subunit expression, seems to suggest an excitatory/inhibitory (E/I) imbalance, operating through multiple mechanisms and converging to produce circuit hyperexcitability. In fact, persistent elevation of GluA1 and GRIN1 subunits suggests enhanced glutamatergic transmission in spinal nociceptive circuits and appear in line with similar data in the literature indicating that offspring of high LG-ABN mothers display increased expression of the NMDA receptor subunit in the hippocampus (Liu et al., 2000). As a key subunit of NMDA receptors, GRIN1 transcripts upregulation might also promote NMDA receptor-mediated transmission and calcium permeability. AMPA receptors are traditionally divided into two functionally distinct groups based on the inclusion or exclusion of GluA2, a subunit determining calcium permeability (Geiger et al., 1995; Isaac et al., 2007; Miguez-Cabello et al., 2025). Thus, AMPA receptors lacking edited GluA2 are calcium permeable and of higher conductance (Isaac et al., 2007). This subunit defect has recently been proposed to be a cause of neurodevelopmental disorders (Salpietro et al., 2019). Additionally, higher GluA1/GluA2 mRNA ratio is directly associated with increased calcium permeable AMPA receptors (CP-AMPARs) and has been suggested to serve as a molecular switch in the formation of GluA2 lacking CP-AMPARs (Pellegrini-Giampietro et al., 1997; Kondo et al., 2000; Guo and Ma, 2021). In our study, the GluA1 transcript global upregulation in the SC, as well as the decrease in GluA2 observed in NMS DRG at P24 could therefore further modify the system toward calcium-permeable AMPA receptors and increased excitability. This molecular configuration would also create vulnerability to excitotoxicity and promote central sensitization (Guo and Ma, 2021), which are cellular mechanisms classically underlying pain hypersensitivity and chronic pain conditions (Yezierski et al., 1998; Fitzgerald, 2005; Longoni and Ferrarese, 2006). On the other hand, the specific GABAA receptor subunit profile observed in the SC from NMS rats provides critical mechanistic insight into why increased maternal care failed to confer resilience. In fact, natural high LG-ABN increases α1- and γ2-containing GABAA subunits in amygdala and hippocampus, enhancing GABAergic synaptic transmission and stress resilience; while low LG-ABN offspring showed increased levels of α3 and α4 subunit-containing GABAA receptor mRNAs (Caldji et al., 2003). Here, GABAAα2, the dominant subunit mediating spinal analgesia through benzodiazepine-sensitive receptors (Knabl et al., 2008; Witschi et al., 2011; Ralvenius et al., 2015), show transient upregulation in the DRG of NMS offspring at P14 but remain unchanged in the spinal cord at both timepoints. NMS also induces persistent spinal increases in α1 and α4 subunits. This profile, characterized by α1 and α4-containing GABAA receptors elevation without α2 enhancement, differs from adaptive GABAergic responses and points out a stress-reactive rather than analgesic receptor composition. In particular, α1-containing receptors are mostly concentrated around the central canal of the spinal cord, while nociceptive afferent terminate in the most superficial layers of the dorsal horn, where α2 is present (Zeilhofer et al., 2009; Ralvenius et al., 2015). Moreover, α4-containing GABAA receptors are benzodiazepine-insensitive (Yang et al., 1995; Sigel and Ernst, 2018), predominantly extrasynaptic and mediate tonic inhibition (Farrant and Nusser, 2005; Lorenz-Guertin and Jacob, 2018), differing functionally from the synaptic phasic inhibition mediated by α1 /γ2 receptors. On the other hand, the α4 subunit undergoes marked changes in expression as a response to fluctuating levels of neurosteroids, which have been shown to be crucial for the processing of nociceptive information at the spinal cord level during development and inflammatory states (Keller et al., 2004; Poisbeau et al., 2005; Smith et al., 2007). Confirming our results, ELS studies on GABAA receptors already hinted that ELS produces long-lasting changes in benzodiazepine binding site number and alterations in GABAA receptor subunit expression (Skilbeck et al., 2010).

To conclude, rather than representing a maternal neglect model, our findings align more closely with emerging models of fragmented maternal care (Baram et al., 2012; Glynn and Baram, 2019). In both rodent models and human observational studies, fragmented patterns of maternal behaviors have been identified as critical for offspring neurodevelopment, predicting hippocampal dysfunction, impaired memory, and elevated risk for anxiety and depressive-like behaviors (Baram et al., 2012; Davis et al., 2017; Aran et al., 2024). In this study, NMS presents an increased reactive maternal care, temporally fragmented by repeated separation-reunion cycles, and potentially delivered in an anxious physiological state. The resulting phenotype of this research (neurotrophic dysregulation, disrupted chloride homeostasis, E/I imbalance, and nociceptive hypersensitivity) could extend previous findings suggesting that fragmented maternal care patterns can also affect peripheral and central somatosensory systems, with long-term consequences for pain processing. In human populations, identifying and supporting at-risk mother-infant dyads during critical postnatal windows (with interventions targeting not just the quantity but the consistency and predictability of maternal care) may help to mitigate long-term neurodevelopmental vulnerabilities, including those affecting pain sensitivity.

## Acknowledgments

Research work has been financed by recurrent funding from the CNRS and the University of Strasbourg, as well as by grants from the Agence nationale de la recherche (ANR) as part of the Programme d’investissement d’Avenir (ANR-17-EURE-022 contract, EURIDOL), the Région grand est (ClueDOL, Fonds de coopération régionale et de recherche). PP is a senior member of the Institut Universitaire de France. HI received a fellowship from french Ministère chargé de l’enseignement supérieur et de la recherche. The authors thank Dr. Pierre-Eric Juif for previous work on the NMS model.

