## Supplemental Table 1 and Figures 1-4 for "Maternal behavioral compensation after neonatal separation fails to prevent spinal circuit reprogramming in offspring"

**Supplementary data**

| **Gene** | **Primers sequences (forward/reverse)** | **NCBI reference** |
| --- | --- | --- |
| **HPRT** | TGGTGAAAAGGACCTCTCGAA  TCAAGGGCATATCCAACAACA | NM013556.2 |
| **GluA1** | CTGGTTGCCTTAATCGAGTTC  GGCTTCATTGATGGATTGCT | NM031608 |
| **GluA2** | CAAGAAGCCTCAGAAGTCCAA  CCCAATGTAGGCAAACACAAT | NM017261 |
| **GRIN1** | CATCACGGGCATCAATGA  CCACCTGCCTCCGGAAGTAGA | NM017010 |
| **GABA_A_ α1** | GACCGTTCTGACCATGACAAC  CACGAAGGCATAGCACACTG | NM183326 |
| **GABA_A_ α2** | TCAGACCTATCTGCCTTGCAT  ACTCCAAACACAGTTCTCGCT | NM001135779 |
| **GABA_A_ α4** | CCATGAGACTGGTGGATTTT  CTTTGGTCCAGGTGTAGATCA | NM080587 |
| **GABA_A_ γ2** | AGGTCATGGAGGCTGTATCA  TCAGATCAAAGTACACGGACA | NM183327 |
| **GABA_A_ β2** | GGGTCTCCTTTTGGATCAACT  TGGGTATTGATTGTGGTCATC | NM012957 |
| **BDNF** | CACAGTCCTGGAGAAAGTCCC  CCTTCCTTCGTGTAACCCAT | NM012513 |
| **GDNF** | GTGTTGCTCCACACCGCGTCT  GGTCTTCAGCGGGCGCTTC | BC119031 |
| **NT3** | AAGTCCTCAGCCATTGACATT  GCTTCTTTACACCTCGTTTCA | NM031073 |
| **NT4** | CCTGCTCTCTCCTCCTTTTCC  AGGTCCCACTCAGGAGCCAGA | NM013184 |
| **NKCC1** | GGGCCTCCTCACACGAAGAA  TGAGGAGCCGAGGGTACTTCA | NM019229 |
| **KCC2** | ACTACAGCTGGCCACCTCGC  ATGCTGCCCTCAGAGAAACGC | NM134363 |

Supplementary Table 1: Primer sequences used in this study.

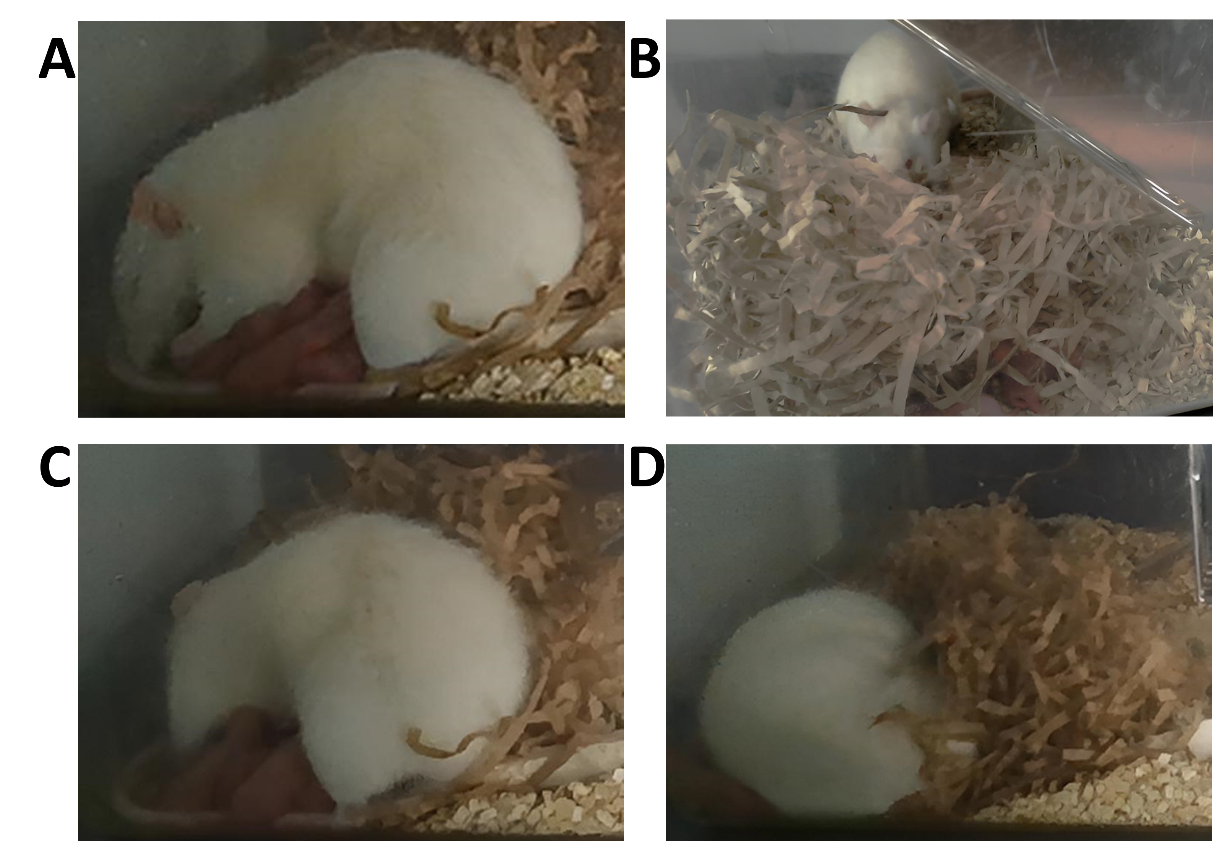

Supplementary Figure 1: Pictures of manifestations of caring maternal behaviors. A: Licking/grooming, B: Nest building, C: Active nursing, D: Passive nursing. Each dam was observed for 30min and behaviors were timed before or after pup retrieval test (control group) or NMS.

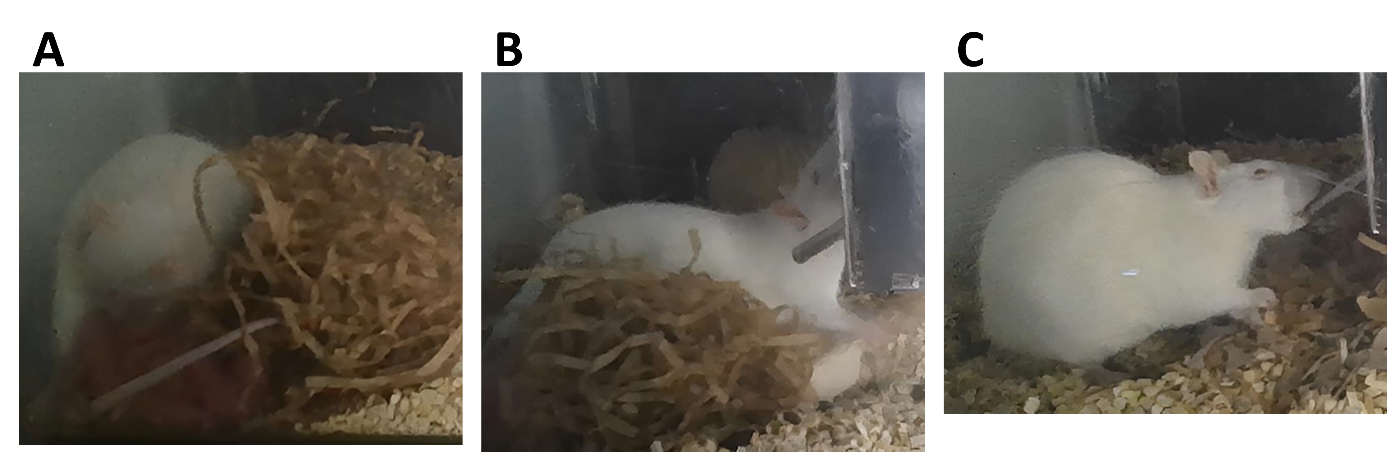

Supplementary Figure 2: Pictures of manifestations of self-maintenance behaviors. A: self-grooming, B: Eating, C: Drinking. Each dam was observed for 30min and behaviors were timed before or after pup retrieval test (control group) or NMS.

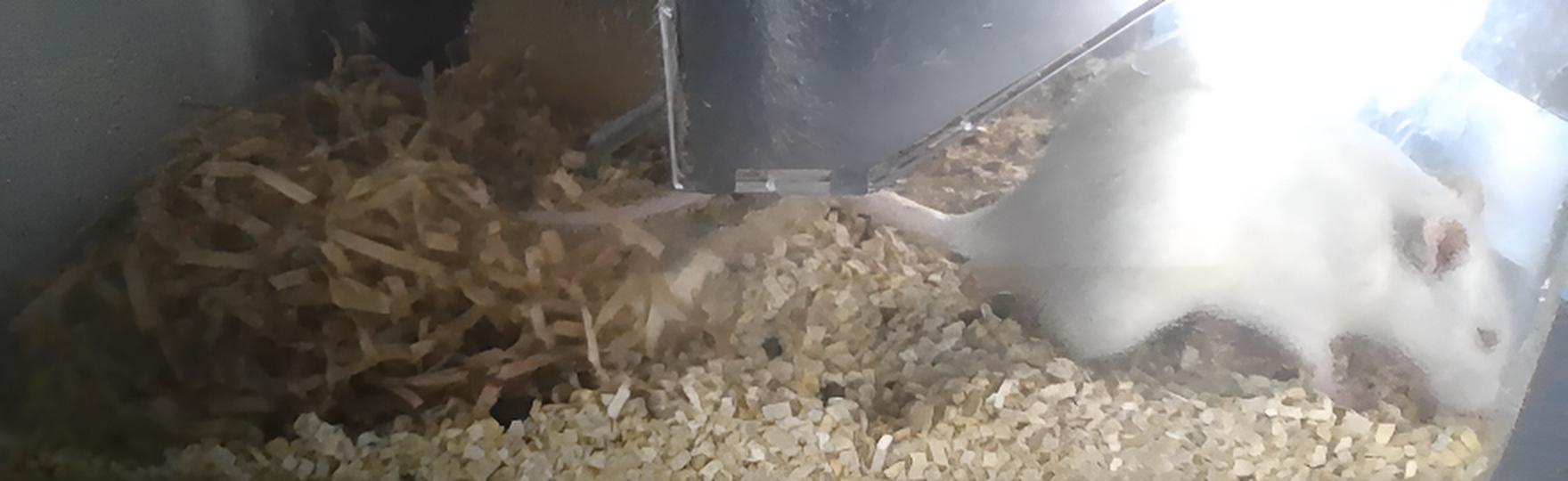

Supplementary Figure 3: Picture of a dam out of nest. Each dam was observed for 30min and behaviors were timed before or after pup retrieval test (control group) or NMS.

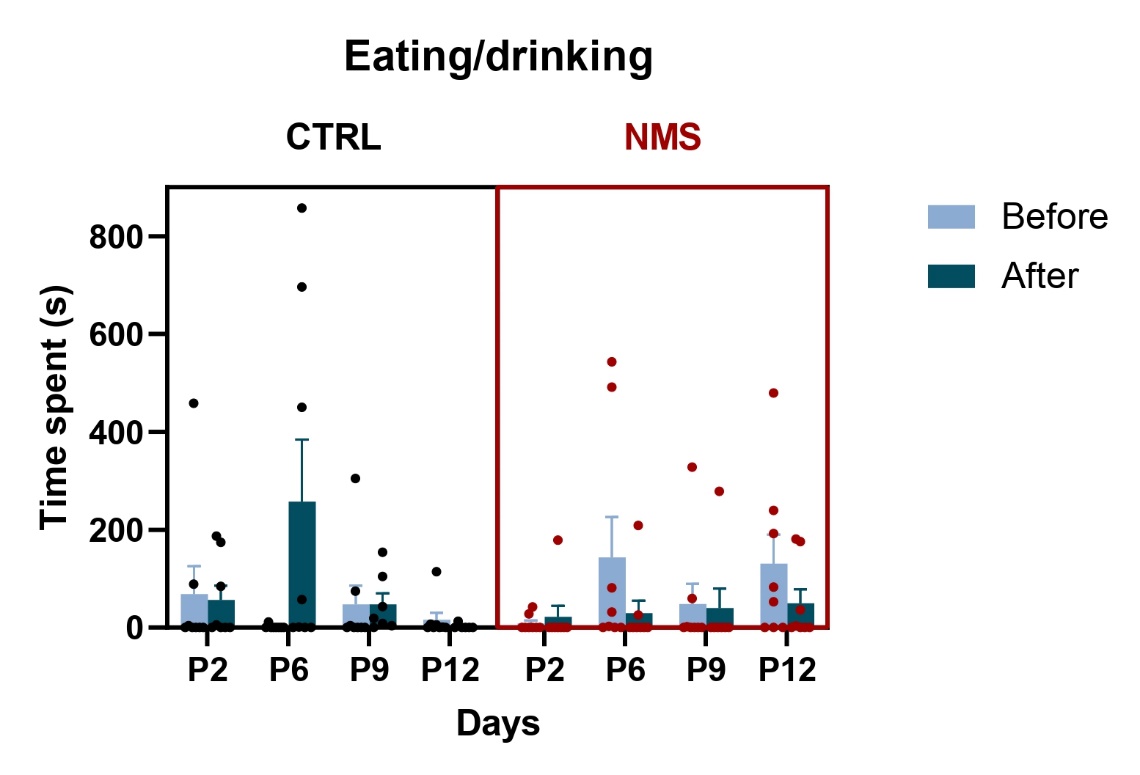

Supplementary Figure 4: Effects of NMS on the range and time allocation of eating and drinking. Each dam was observed for 30min and 7 behaviors were timed before or after pup retrieval test (control group) or NMS. Statistical analysis was performed using LMM on square-root-transformed data. Statistical significance was assessed with 3wANOVA, followed by post-hoc pairwise comparisons using emmeans with Tukey adjustment for multiple comparisons. An **interaction effect of GroupxTiming** was revealed. CTRL: N = 8, NMS: N = 8.
